# Multinuclear MRI reveals early efficacy of stem cell therapy in stroke

**DOI:** 10.1101/2022.02.08.479630

**Authors:** Shannon Helsper, Xuegang Yuan, F. Andrew Bagdasarian, Jacob Athey, Yan Li, Cesario V. Borlongan, Samuel C. Grant

## Abstract

Compromised adult human mesenchymal stem cells (hMSC) can impair cell therapy efficacy and further reverse ischemic recovery. However, *in vitro* assays require extended passage to characterize cells, limiting rapid assessment for therapeutic potency. Multinuclear magnetic resonance imaging and spectroscopy (MRI/S) provides near real-time feedback on disease progression and tissue recovery. Applied to ischemic stroke, ^23^Na MRI evaluates treatment efficacy within 24 h after middle cerebral artery occlusion, showing recovery of sodium homeostasis and lesion reduction in specimens treated with hMSC while ^1^H MRS identifies reduction in lactate levels. This combined metric was confirmed by evaluating treatment groups receiving healthy or compromised hMSC versus control (sham saline injection) over 21 d. Behavioral tests to assess functional recovery and cell analysis for immunomodulatory and macrophage activity to detect hMSC potency confirm MR findings. Clinically, these MR metrics may prove critical to early evaluations of therapeutic efficacy and overall stroke recovery.

## Introduction

Cell therapies, particularly human bone-marrow derived mesenchymal stem cells (hMSC), are the focus of clinical trials and preclinical research for the treatment of neurological diseases [1]. To date, 29 studies worldwide involve hMSC in the treatment of stroke (ClinicalTrials.gov). Clinical use of hMSC is desirable due to their homing ability to injury, secretion of soluble factors critical for cell survival and proliferation, and modulation of immune responses [2,3]. In addition to paracrine and immunomodulatory functions, hMSC-based therapies offer low immunogenicity, easy isolation and multiplication, making them ideal therapeutic agents [4]. Enhanced by culture conditions, the natural release of neuroprotective, anti-inflammatory, vasogenic and neurotrophic factors [5,6] is particularly advantageous for treatment of neurological diseases [2], such as ischemic stroke [7].

Therapeutic efficacy of hMSC is dependent on multiple parameters, including culture-based preconditioning, *in vivo* stem cell response and donor heterogeneity [8]. Extended culture and preconditioning metrics should be monitored to ensure that high therapeutic efficacy is maintained. In this fashion, extensive *in vitro* assays may predict *in vivo* stem cell performance in clinical applications [2]. At present, however, extended passage for *in vitro* characterization to assess therapeutic potential of hMSC reliably beyond the point of administration is not common practice. In 2005, the first clinical trial to investigate the safety and efficacy of human stem cells performed minimal *in vitro* testing prior to patient administration [9]. Simple surface expression markers for SH-2 and SH-4 were measured via flow cytometry, while cell viability and culture sterility were checked. Robust *in vitro* analysis and investigation into therapeutic variance is imperative to assist translational studies of hMSC therapies to minimize deficits arising from cell or source variance [8].

While *in vitro* cell assays can assist to predict therapeutic efficacy, magnetic resonance imaging and spectroscopy (MRI/S) enable real time monitoring of the *in vivo* efficacy and evolution of hMSC therapies, starting at therapeutic administration and following through disease progression and recovery. MRI/S offer multiple advantages over other imaging modalities, including noninvasive monitoring of functional tissue recovery as well as assessment of implanted cell biodistribution [10]. Using internalized super-paramagnetic iron oxide (SPIO) particles to generate strong negative contrast, initial delivery and migration of labeled hMSC can be monitored [10,11], in concert with longitudinal cell detection at high resolution [12,13]. In the case of stroke treatment, MRI/S can provide functional information of recovery at the neuronal level in addition to hMSC tracking.

At present, the ‘golden standard’ for acute stroke imaging is ^1^H diffusion-weighted MRI due to its noninvasive detection of an ischemic event within minutes of onset [14]. However, diffusion-weighted imaging and more recent diffusion-perfusion mismatch techniques are limited in their ability to assign accurately and consistently core volumes versus penumbra, especially in longitudinal studies [15]. Sodium MRI overcomes this limitation, providing functional recovery information at the cellular level. Transmembrane Na^+^ and K^+^ gradients play a key role in maintaining the cellular resting membrane potential, osmotic pressure and cell volume [16]. These processes are highly sensitive to ATP depletion, consuming up to one third of the cell’s total energy [16]. Under ischemia, namely low O_2_ and ATP, maintenance of the Na^+^/K^+^ -ATPase pump is disrupted, and a detrimental cascade of neurological events occurs, including an influx of sodium into the cell. Taking advantage of the enhanced quadrapolar MRI sensitivity at high field, acute sodium influx becomes measurable, and ^23^Na MRI can provide an important quantitative metric to assess ischemia and homeostatic recovery directly [15,17]. Likewise, Magnetic Resonance Spectroscopy (MRS) also offers a functional investigation of ischemic tissue evolution via acute metabolic activity. The recently developed relaxation-enhanced MRS (RE-MRS) provides quantifiable biochemical measurements of cell metabolites, using highly selective spectral excitation to target resonances of interest exclusively and avoiding the influence of water. In stroke models, RE-MRS provides access to key metabolites of interest: choline (Cho) for membrane synthesis, creatine (Cre) and lactate (Lac) with respect to cellular bioenergetics and N-acetyl aspartate (NAA) as an indicator of compartment-specific viability [18–20]. At present, both ^23^Na MRI and ^1^H MRS remain underutilized in clinical stroke assessment, in particular with respect to treatment evaluation.

The present study investigates the utilization of MRI/S to assess treatment efficacy of hMSC in a stroke model and its clinical potential to complement current diffusion-weighted MRI. Early cell passage *in vitro* analysis corresponding to the time of implantation indicated comparable therapeutic potential between two sets of adult hMSC. Robust *in vitro* analysis for extended passages confirmed that one set of cells was ‘compromised’ while the other remained ‘healthy.’ The impact of this therapeutic variance on tissue recovery was evaluated with 3D ^23^Na chemical shift imaging (CSI) and ^1^H RE-MRS in conjunction with and compared to traditional ^1^H T_2_W MRI in a rat transient middle cerebral artery occlusion (MCAO) model of ischemic stroke.

## Materials and Methods

### Cell Culture and 3D aggregation

Both hMSC lines (compromised and healthy) from passage 0 to 2 (P0-2) were acquired from the Tulane Center for Gene Therapy. hMSC were isolated from the bone marrow of donors aged 19-49 y based on plastic adherence. Both cell lines were negative for CD3, CD14, CD31, CD45, CD117 (all less than 2%), positive for CD73, CD90, CD105 and CD147 markers (all greater than 95%) and possess tri-lineage differentiation potential upon *in vitro* induction. Frozen hMSC suspensions containing 10^6^ cells/mL/vial in freezing media that contained α-MEM, 2-mM L-glutamine, 30% fetal bovine serum (FBS) and 5% dimethyl sulfoxide (DMSO), were thawed and cultured [7,21]. In brief, hMSC were expanded and maintained in complete culture media (CCM) containing α-MEM with 10% FBS (Atlanta Biologicals, Lawrenceville, GA) and 1% Penicillin/Streptomycin (Life Technologies, Carlsbad, CA), exchanging the medium every three days. Cells were grown to 70-80% confluence and then harvested by incubation with 0.25% trypsin/ethylenediaminetetraacetic acid (EDTA) (Invitrogen, Grand Island, NY). Harvested cells were re-plated at a density of 1,500 cells/cm^2^ and sub-cultured up to P8. Cells at P5 and P8 were used for aggregate preconditioning, *in vitro* cellular assessment and transplantation. Preconditioning of cells prior to *in vivo* administration is a critical step to regulate and enhance therapeutic potential. Aggregate preconditioning of cells has been used to mimic a three dimensional, low oxygen micro-environment, which in turn improves viability under ischemia [22–24]. Increased cell compaction from aggregation also enhances cell-to-cell interaction, ultimately contributing to the increased cytokine secretion [21] and enhanced cell migration [25].

3D hMSC aggregates were obtained by resuspending harvested hMSC in CCM, adding 200,000 cells into each well of ultra-low attachment (ULA) 6-well culture plates (Corning, Corning, NY). To induce spontaneous aggregate formation, ULA plates were placed on a programmable rocking platform with 8° rocking angle and 20 RPM in a standard CO_2_ incubator (37 °C and 5% CO_2_) for 48 h. hMSC aggregates were dissociated with 0.25% trypsin/EDTA (Invitrogen) to acquire single cell suspension. hMSC obtained from dissociated aggregates were cultured subsequently on tissue culture surfaces for recovery over two days under 37°C and 5% CO_2_. Recovered hMSC were defined as aggregate-derived dissociated hMSC. Before transplantation, hMSC were incubated with MPIO (7.47-μg Fe/mL) of for 12 h. Subsequent rinsing and re-suspension in phosphate saline buffer was done just prior to injection. All reagents were obtained from Sigma Aldrich (St. Louis, MO) unless otherwise noted.

### Cell Number, CFU-F, SA-β-Gal Activity and immunomodulation

Cell numbers were determined with fluorescence spectroscopy using a Quant-iT^™^ PicoGreen kit (Invitrogen, Grand Island, NY). In brief, cells were harvested, lysed over-night using proteinase K (VWR, Radnor, PA), and stained with Picogreen to measure cellular DNA content. Fluorescence signals were read using a Fluror Count (PerkinElmer, Boston, MA). Population doubling time (mean PD time) was determined through culture in each passage:

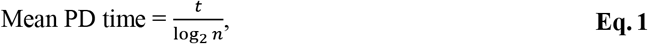

where n is the cell number fold increase during culture time, t.

For CFU-F assays, hMSC were harvested and re-plated at 15 cells/cm^2^ on a 60-cm^2^ culture dish and cultured for another 14 d in CCM. Cells were stained subsequently with a solution containing 20% crystal violet in methanol for 15 min at room temperature. Stained cells were rinsed three times with PBS. The number of individual colonies were counted manually. Cellular senescence was evaluated by SA-β-Gal activity assay kit (Sigma, St. Louis, MO) as described in the manufacturer’s instructions. To evaluate the immunomodulatory potential of hMSC, indoleamine 2,3-dioxygenase (IDO) enzymatic activity (including both IDO1 and IDO2) was determined via Kynurenine level in culture medium. A 400-μL cell culture medium was clarified by addition of trichloroacetic acid (200-μL, 30% by weight; Sigma Aldrich, St. Louis, MO), followed by vortex mixing and centrifugation at 8,000g for 5 min. An equal volume of Ehrlich reagent (2% p-dimethylaminobenzaldehyde in glacial acetic acid) was added to the clarified supernatant, and optical density at 490 nm was measured via Fluror Count (PerkinElmer, Boston, MA). The optical density was normalized to sample cell number.

### Real-time Reverse Transcriptase-Polymerase Chain Reaction

Total RNA was isolated using the RNeasy Plus kit (Qiagen) following manufacturer’s instructions. Reverse transcription was carried out using 2 μg of total RNA, anchored oligo-dT primers (Operon) and Superscript III (Invitrogen). Primers for specific target genes were designed using the software Oligo Explorer 1.2 (Genelink). β-actin was used as an endogenous control for normalization. RT-PCR reactions were performed on an ABI7500 instrument (Applied Biosystems) using SYBR Green PCR Master Mix. The amplification reactions were performed, and the quality and primer specificity were verified. Fold variations in gene expressions were quantified using the comparative Ct method: 2^-(CtTreatment-CtControl)^, which is based on the comparison of the target gene (normalized to GAPDH) among different conditions.

### Macrophage differentiation and co-culture with hMSC

Human monocyte line THP-1 was purchased from ATCC and used to establish human naïve macrophage differentiation and polarization. Briefly, THP-1 cells were grown in RPMI1640 (ATCC) containing 10% FBS, 1% Penicillin/Streptomycin and 0.05-mM 2-mercaptoethanol. For M0 differentiation, THP-1 cells were treated four days with 20-nM Phorbol 12-myristate 13-acetate (PMA) for a complete M0 phenotype. To generate M1 macrophages, M0 macrophages were polarized with 20-nM PMA, 20-ng/mL recombinant human IFN-γ (Peprotech, Rocky Hill, NJ), and 100-ng/mL lipopolysaccharide (LPS) for two days to induce the inflammatory phenotype. To generate M2 macrophages, M0 macrophages were polarized with 20-nM PMA and 10-ng/mL IL-4 (Peprotech, Rocky Hill, NJ) for two days to induce an anti-inflammatory phenotype. To investigate the influence of hMSC from different donors on the differentiation from monocyte to M0 macrophage, both cell types were co-cultured in a six-well transwell plate (Corning, Corning, NY). M0 macrophage was induced at the bottom of a six-well plate for two days, and then hMSC were added on the upper culture insert at a ratio of hMSC:THP-1=1:10. Co-culture was maintained under M0 differentiation conditions for another two days, and M0 macrophages were collected for RT-PCR analysis. Similarly, to investigate the influence of hMSC on the M1 or M2 polarization, M1 or M2 polarization was induced at the bottom of six-well plate along with hMSC cultured on the insert. The co-culture was maintained under M1 or M2 polarization for two days, and the macrophages were collected for RT-PCR analysis.

### *In vitro* OGD model and iNC-hMSC co-culture

iNC were differentiated from human induce pluripotent stem cells (iPSC) as described in previous studies and allowed to mature for seven days. Subsequently, the oxygen-glucose deprivation (OGD) model [26] was induced as follows: neural culture medium was replaced with a glucose-free DMEM medium suppled with 1% Penicillin/Streptomycin. Cells were cultured in a humidified oxygen-controlled C-chamber (BioSpherix, Lacona, NY) in a standard cell culture incubator at 37°C, with continuous flow of 94.8% N2 and 5% CO_2_ gas. OGD was carried out for 90 min and then terminated by returning the culture back to standard 20% O_2_ and 5% CO_2_ culture incubator with neural culture medium for one hour of reperfusion. 20,000 hMSC were seeded on each insert of a 12-well transwell culture plate then placed on top of OGD-treated iNC. The co-culture was maintained for four hours, and the iNC were collected for Live/Dead staining (ThermoFisher) via flowcytometry analysis as manufacture’s instruction. An exact gating strategy was applied to all samples.

### Animal Model

All animal procedures were completed in accordance with the Animal Care and Use Committee at the Florida State University. A transient middle cerebral artery occlusion (MCAO) [27] simulating striatal ischemia was induced in Sprague-Dawley rats (200–250 g, RRID 5651135) by surgically introducing a rubber-coated filament in the circle of Willis for one hour. In brief, rats were anesthetized with 4-5% isoflurane in 100% O_2_ and maintained at ~3% anesthesia for the duration of surgery. A rubber-coated filament (Doccol Corp., Sharon, MA) was inserted into the external carotid artery and threaded 1.9-cm into the internal carotid artery to block the circle of Willis, effectively occluding the MCA. The filament was secured and midline incision was sutured temporarily while the rats were allowed to recover in a warmed incubator for the duration of the one-hour occlusion. Rats were re-anesthetized for filament removal and immediate arterial injection of ~1 mil hMSC from compromised (n=7) or healthy donors (n=9) in 50-μL saline or saline only (control, n=11). All received pre- and post-operative analgesics (bupivacaine and buprenorphine, respectively) as well as saline for rehydration, with additional administration the next day after the first MR session. Baseline behavioral assessments were conducted with subsequent testing concurrent with MR scanning to 21 d post-ischemia.

### Magnetic Resonance Acquisitions

All MR data was acquired using the 900-MHz vertical magnet (21.1-T) at the National High Magnetic Field Laboratory in Tallahassee, FL [28]. The spectrometer was equipped with a Bruker Avance III console operated using Paravision 5.1 (Resonance Research, Inc. Billerica, MA). A home-built ^23^Na/^1^H double-tuned linear birdcage coil was used to image the animals, enabling to evaluate tissue injury and recovery. Rats were secured in a custom cradle and maintained under anesthesia at or below 3% isoflurane. To further minimize motion during acquisitions, respiratory triggering (Small Animal Instruments, Inc., NY) was used. To assess tissue recovery and treatment efficacy, MR imaging was completed at 1-, 3-, 7- and 21-d post-ischemia. Localization images were acquired using a ^1^H Rapid Acquisition with Relaxation Enhancement (RARE) pulse sequence. Cell administration and subsequent clearing was confirmed with a 2D gradient recalled echo (GRE) sequence at 50×50-μm in-plane resolution [TE = 4 ms, TR = 1 s]. T_2_-weighted images were generated utilizing a ^1^H 2D FSE sequence with 100×100-μm in-plane resolution [TE = 12.5 ms (effective 25 ms), TR = 6 s]. 3D ^23^Na chemical shift imaging (CSI) was acquired at 1-mm isotropic resolution [TR = 60 ms], with a total acquisition time of 32 min. Relaxation-enhanced MR spectroscopy (RE-MRS) enabled evaluation of metabolite concentrations in both the ischemic and contralateral hemispheres. Selective bandwidth excitation pulses targeted lactate (Lac), creatine (Cre), choline (Cho) and N-acetyl aspartate (NAA), while avoiding water. Spatial selectivity was introduced by adiabatic selective refocusing (LASER) [18–20] pulses. T_2_W images enabled anatomical reference to the ischemic lesion and contralateral alignment for the localized voxels.

### Analysis

CSI data reconstructed in MATLAB was zero-filled to 0.5-mm isotropic resolution for volumetric and signal analysis in Amira 3D Visualization Software (Thermo Fisher Scientific, Waltham, MA). A signal threshold generated from the contralateral hemisphere was used to define the ischemic lesion:

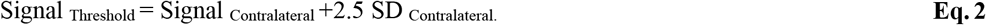

All signal above this threshold, excluding CSF, was assigned to the ischemic lesion. ^1^H T_2_W data were reconstructed in Amira in the same manner. RE-MRS data were reconstructed in JMRUI using Linear Prediction Singular Value Decomposition to select components with peaks assigned according to previous literature, NAA 2.0 ppm, Lac 1.31 ppm, Cre 3.0 ppm, Cho 3.2 ppm [29], referencing water at 4.7 ppm. Metabolite ratios to choline were used to track metabolic signals longitudinally. Statistical analysis was completed using JMP 14.0 (SAS Institute Inc., Cary, NC) for MR data and GraphPad Prism 9.0.2 (GraphPad Software, San Diego, CA) for cell assays. Statistical significance is shown according to a mixed model with pairwise Student T test for post-hoc comparison for MR data and unpaired T test for cell assays. Outliers in MR data, defined as greater than 1.5 times the interquartile range, were excluded. All graphs are mean ± standard deviation.

### Behavioral and Neuromotor Assessment

All animals underwent baseline behavior assessment prior to surgery. Assessments were done in the same order with at least ten minutes of rest in the home cage between each test. Temperature, lighting and noise were monitored while personnel access during testing was restricted. Test recordings enabled the video reviewer to be blinded to treatment groups. For the open field test, rats were placed in the center of the 92×92×56-cm grid and equally distributed 36 square grids for ten minutes. Total distance traveled and time spent within the grid center compared to exterior was determined. A spontaneous forelimb usage asymmetry score was assigned using the cylinder test, as previously described [30,31]. In brief, rats were placed in a tall transparent vertical cylinder for ten minutes and the number of independent forepaw contacts with the cylinder surface were recorded. In addition, the elevated body swing test [32] consisted of twenty swings in which rats were lifted vertically above a table by the tail approximately 15 cm and immediately returned to resting position on the table following each swing. Directional preference for each swing were recorded.

## Results

### Combined multinuclear [^23^Na, ^1^H] MR/S detects successful therapy 24 h after administration

To confirm healthy hMSC and differentiate from control (vehicle only) or ‘compromised’ therapy applied to ischemic stroke, a multinuclear MR approach to quantify cerebral sodium homeostasis and lactate levels was developed. 3D ^23^Na CSI was applied to all subjects to monitor the ischemic lesion progression both in terms of volume and sodium signal to compare between hemispheres and treatment groups. Relaxation-enhanced MRS (RE-MRS) was applied within the ischemic lesion and corresponding contralateral hemisphere to determine metabolic changes, with specific focus on lactate progression. Applied adult hMSC were characterized as ‘healthy’ or ‘compromised’ following extensive *in vitro* analysis and stress tests that otherwise would not have been differentiable using conventional cell assays at the time and passage of implantation. **Fig. 1a** demonstrates the experimental timeline of this study.

**Fig. 1.**
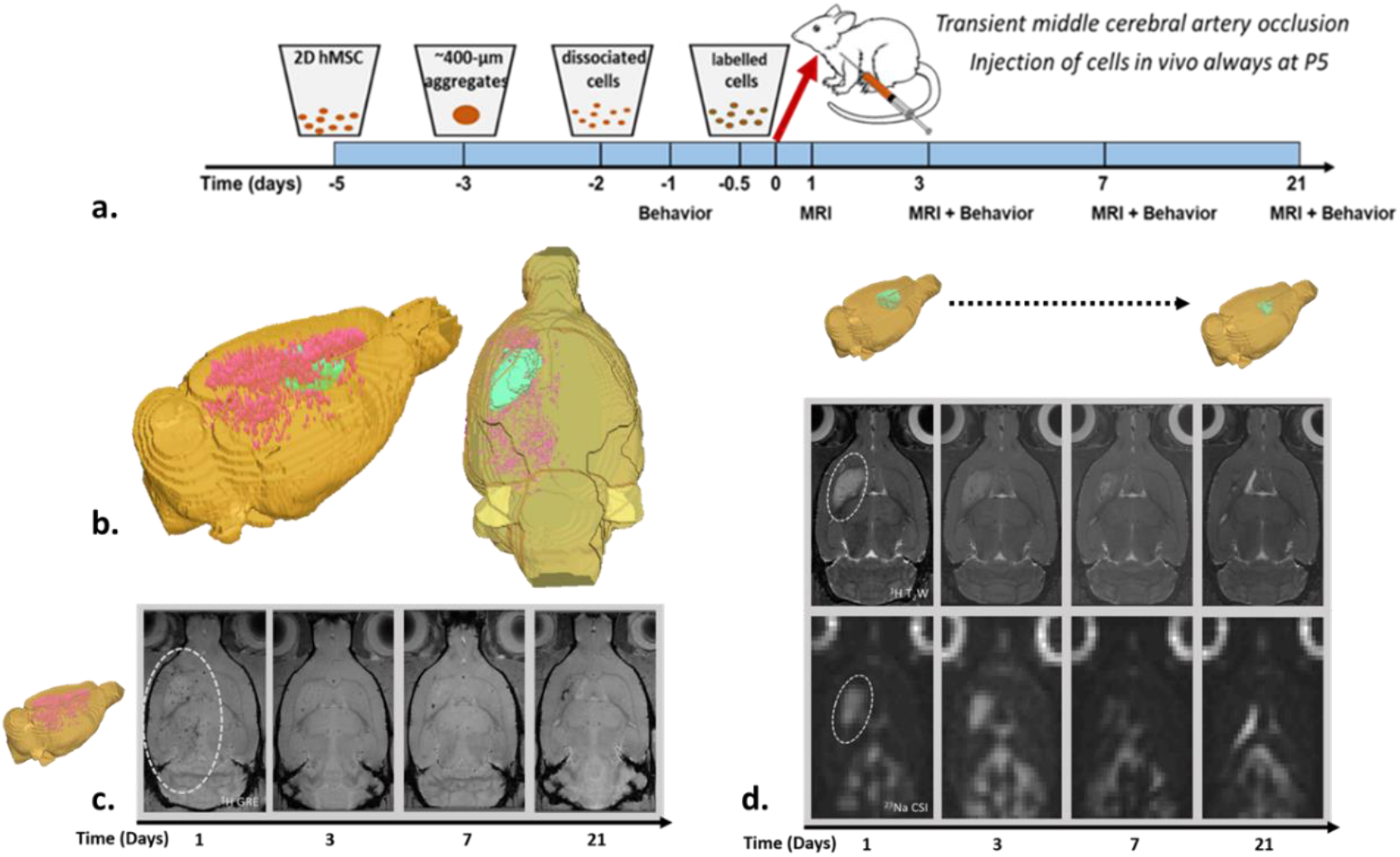
Hyperintense signal in ^23^Na and ^1^H MRI depict ischemic lesion decreasing over time and confirms successful cell administration. a) Experimental timeline. b) 3D rendering of ^1^H MRI rat brain depicting ischemic lesion (green) and cells (red) on day 1. Representative MRI of a rat brain following 1 h MCAO and administration of healthy hMSC over the 21-d time course for c) cellular localization (red segmentation corresponding to cells on day 1), d) ischemic lesion assessment via ^1^H T_2_W and ^23^Na CSI with 3D renderings (green segmentation corresponding to ^1^H).

### Ischemic lesion detection and cell therapy traceability was achieved with MRI

Initial ischemic injury was verified one day after MCAO was instituted using conventional 2D ^1^H T_2_W (**Fig. 1d**). Ischemic lesions were defined as hyperintense signal regions in the brain. The 1-h occlusion resulted in all animals exhibiting an ischemic lesion restricted to the striatum, minimizing initial injury variability from surgery opposed to treatment.

Once successful ischemic induction was confirmed, the animal underwent further MR to verify successful delivery (**Fig. 1c**). MR detection of cellular therapy is key to confirm successful administration. Labeling of hMSC with micron-sized particles of iron oxide (MPIO, Bangs Laboratories, Inc., IN) prior to intra-arterial injection allowed cells to be tracked using MRI, displaying the unique homing capabilities of hMSC [7,13]. The cells selectively targeted the ischemic hemisphere where they remained, with the majority of cell clearance seen by day 3.

Once ischemic lesion and cell detection were confirmed via conventional methods, rats underwent further screenings within the same MR session to correlate sodium homeostasis in the lesion compared to a corresponding region of interest in the contralateral hemisphere as well as baseline levels determined by naïve rats. These correlations were compared to conventional ^1^H T_2_W methods to determine method sensitivity. Similar to ^1^H T_2_W images, the hyperintense region in ^23^Na CSI also corresponded to the ischemic lesion and decreased over time (**Fig. 1d**).

### Sodium homeostasis recovery enabled early detection of successful therapy

3D ^23^Na CSI enables detection of bulk sodium in the system to quantify tissue sodium in ischemic and contralateral hemispheres with high spatial resolution compared to more conventional 3D ^23^Na GRE sequences [33]. The sodium quantified in 3D ^23^Na MRI results from local tissue sodium fluctuations. As blood supply is restricted, there is a pronounced decrease in oxygen and glucose to the affected brain region, resulting in compromised cellular energy metabolism and malfunction of the Na^+^/K^+^-pump as rapid loss of ATP persists [34], as demonstrated in **Fig. 2b**. ^23^Na MRI is able to measure this change in tissue sodium, related to cellular volume and compartmental sodium concentration, through the sub-acute and chronic stages of ischemic stroke (**Fig. 2a**).

**Fig. 2.**
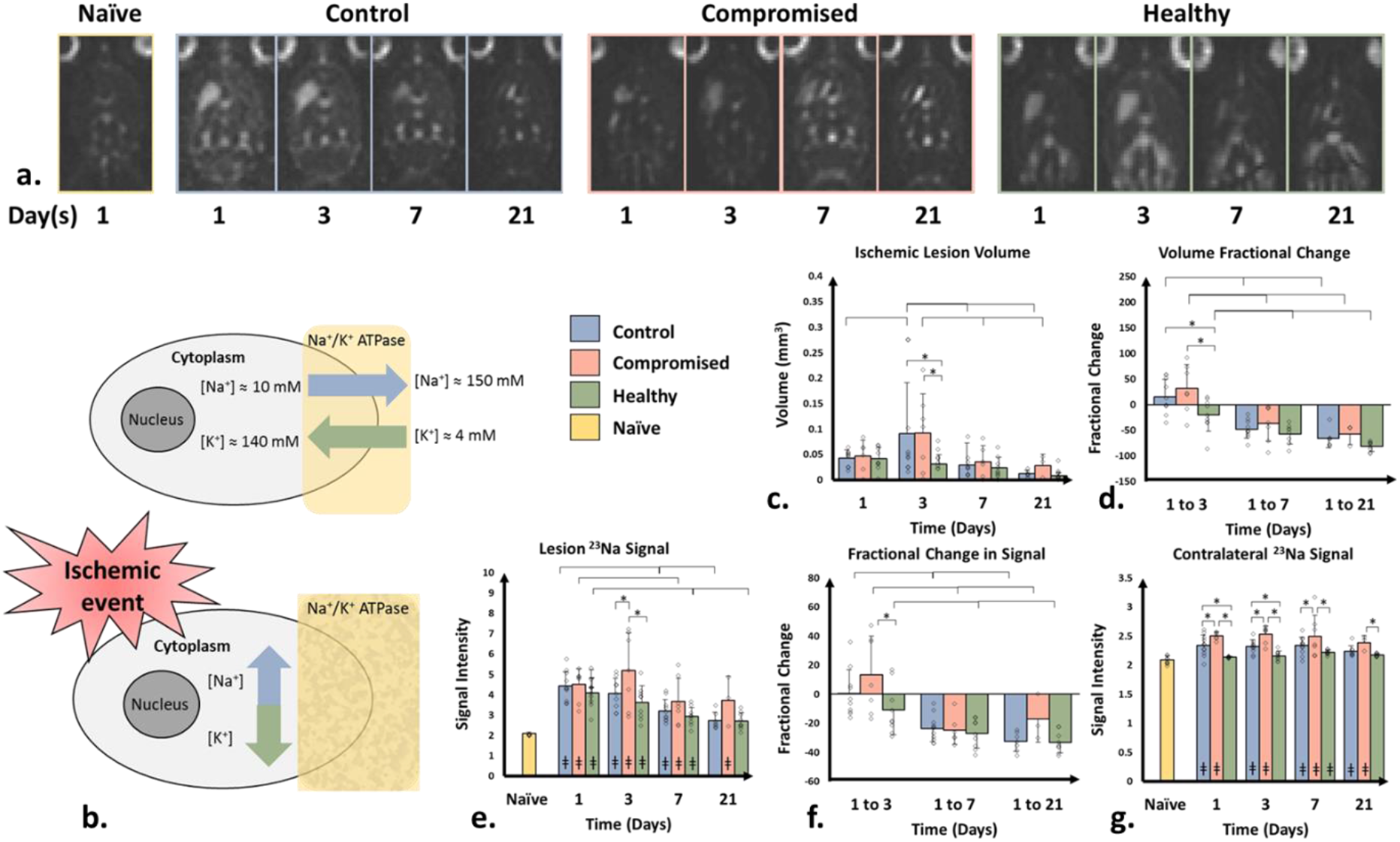
Sodium analysis displays ionic homeostasis recovery in the healthy treatment groupa) 3D ^23^Na CSI 1 to 21 d following MCAO and hMSC administration. b) Schematic of Na^+^/K^+^-ATPase pump. Bar graphs depicting: c) absolute ischemic lesion (Day 3: *, control to healthy p = 0.0.0057; *, compromised to healthy p = 0.0098) d) fractional change in ischemic lesion volume (Day 1 to 3: *, control to healthy p = 0.0187; *, compromised to healthy p = 0.0067), e) ^23^Na signal within region of interest corresponding to ischemic lesion on day 1 superimposed on subsequent days (Day 3: *, healthy to control p = 0.0083; * control to compromised p = 0.0019), f) fractional change in signal (Day 1 to 3: *, p = 0.0218) and g) ^23^Na signal in contralateral (non-ischemic) hemisphere compared between (*) groups (Day 1: control to compromised p = 0.0162; control to healthy p = 0.0022; compromised to healthy p = 0.0001. Day 3: control to compromised p = 0.0012; control to healthy p = 0.0113; compromised to healthy p = 0.0001. Day 7: control to compromised p = 0.0132; compromised to healthy p = 0.0001. Day 21: compromised to healthy p = 0.0205.) and (ǂ) naïve animals. Graphs are depicted as mean with standard deviation error bars. Significance was determined by mixed-model with Student’s T test with p < 0.05.

High-resolution 3D ^23^Na CSI and post-processing enabled segmentation of the ischemic lesion by comparison to the contralateral hemisphere and naïve rats. At 24 h after the induction of ischemia, 3D ^23^Na CSI confirmed a substantial increase in ischemic lesion volume (**Fig. 2c**) and signal intensity (**Fig. 2e**). However, healthy hMSC triggered an immediate and robust decrease in ischemic lesion volume from 24 to 72 h, demonstrating early recovery compared to the control and compromised groups, which were still increasing in volume by day 3. This reduction in lesion volume as impacted by healthy hMSC contrasted the expected evolution in stroke, for which the lesion volume continues to increase, as evident in the control and compromised groups (**Fig. 2d**) [35]. The average ^23^Na signal within the lesion tracks longitudinal recovery, enabling inter-group comparisons. Most notably, when the ischemic lesion volume defined on day 1 was overlaid on subsequent days, the healthy hMSC group exhibited reduced ^23^Na signal within the pre-defined volume, compared to the compromised hMSC group, providing a method to discriminate between viable and efficacious treatments at 72 h after implantation. In strong contrast to healthy hMSC, the compromised group did not recover signal through the entirety of the time course, indicating a lack of significant recovery of Na^+^/K^+^-ATPase activity. The fractional change in signal (**Fig. 2f**) parallels the volume changes, further supporting early recovery by healthy hMSC.

Systemic affects in the brain were assessed by extending analysis into the non-ischemic hemisphere. The contralateral hemisphere exhibited a significant increase in sodium content for the compromised and control groups as early as 24 h after administration (**Fig. 2g**). In contrast, the healthy hMSC group maintained sodium signal comparable to naïve animals, indicating a systemic neuroprotective effect. The control animals did not recover signal until three weeks after MCAO, while the compromised group did not recover at all, supporting the neuroprotective effect induced by effective therapy and potential detriments induced by a compromised stem cell treatment.

Sodium volumetric measurements are an indicator of ionic homeostasis disruption that both the salvageable penumbra as well as the ischemic core. Volume and signal recovery over time indicates reversal of the Na^+^/K^+^-pump dysfunction at the start of an ischemic event, as well as the ability to restore functional membrane potential in the lesion, while maintaining ionic homeostasis in the contralateral hemisphere when an effective therapy is administered, as is the case with the healthy hMSC.

### Traditional proton imaging supports early detection of compromised cells

Edemas that form as a result of cerebral ischemia can be categorized into three groups: cytotoxic, ionic and vasogenic, corresponding respectively to temporal formation [36,37]. Traditional ^1^H T_2_-weighted imaging depicts this water influx, particularly into the intracellular space leading to glial and neuronal swelling from cytotoxic edema as well as that from ionic edema in the dense ischemic tissue [36,37].

By convention, the hyperintense signal in MR images corresponding to the overall edema was assigned to the ischemic lesion. Regions were designated as ischemic lesion using a signal threshold generated from the non-ischemic hemisphere. Total volume and signal intensity were assigned based on this threshold. As such, the ischemic lesion volume from ^1^H T_2_-weighted images provided support of recovery from healthy hMSC administration (**Fig. 3**). Lesion volume was reduced on days 1 and 3 for the healthy compared to the compromised group, as displayed by volume (**Fig. 3c**); however, this metric was not sensitive enough to be seen as a fractional change in volume (**Fig. 3d**). For the ^1^H signal intensity, longitudinal recovery was indicated for all groups (**Fig. 3e**). However, the healthy hMSC group recovered to baseline signal by day 7, compared to day 21. The fractional change in ^1^H signal manifested in a similar trend as volumetric measurements with an immediate decrease in signal for both treatment groups (**Fig. 3f**). The majority of recovery evidently occurred at early time points, as compared to late time points for untreated ischemia. The ^1^H signal of the non-ischemic hemisphere was compared to naïve animals (**Fig. 3g**). Only temporal fluctuations in signal contrast were evident with a reduction in signal indicated for the healthy hMSC group compared to control on day 7.

**Fig. 3.**
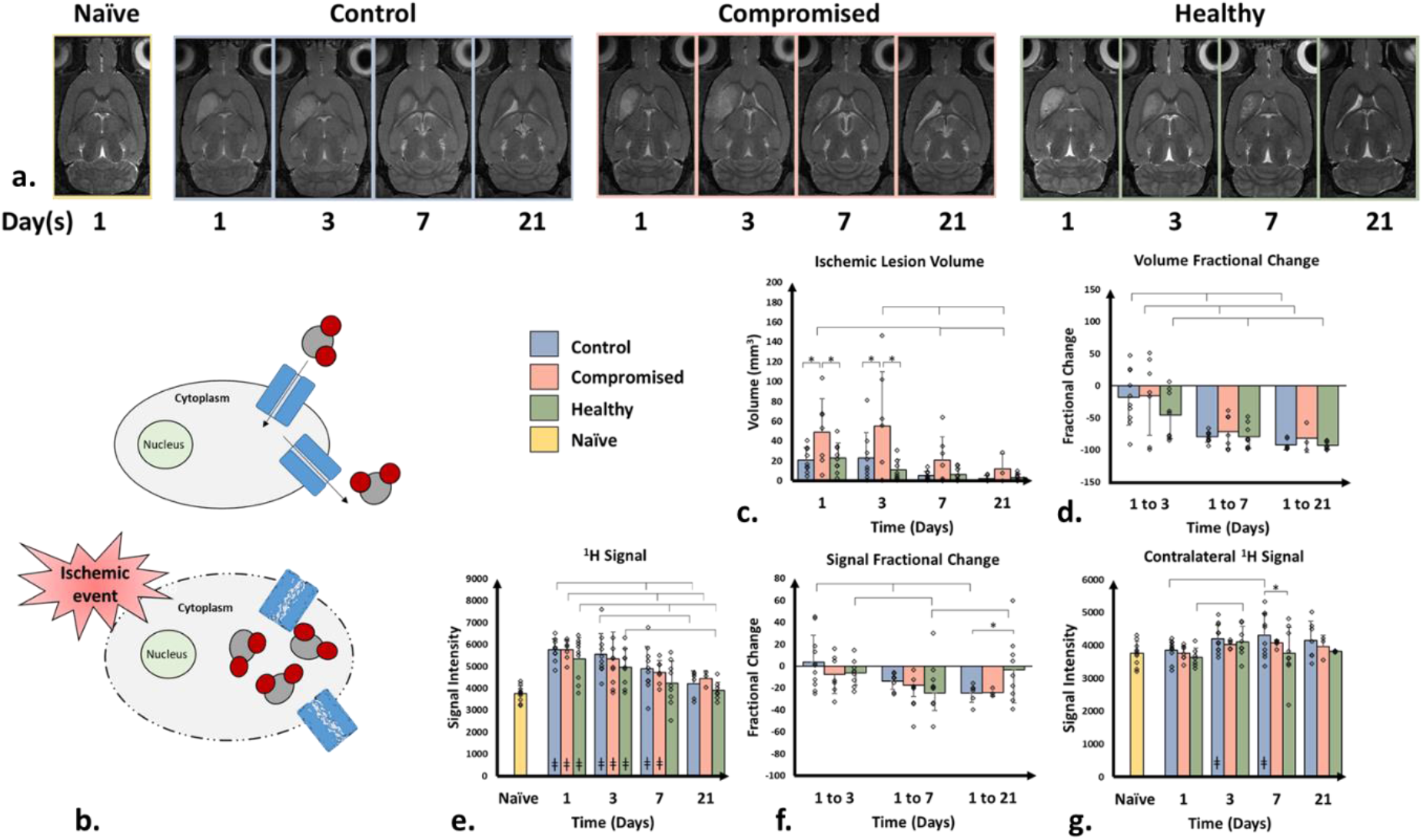
Conventional proton MRI supports sodium assessments in volumetric and signal analysis. a) ^1^H T_2_W MRI 1 to 21 d following MCAO and hMSC administration. Bar graphs depicting: b) Schematic of osmotic swelling c) absolute ischemic lesion (* Day 1: control to compromised p = 0.0122; compromised to healthy p = 0.0202. Day 3: control to compromised p = 0.0042; compromised to healthy p = 0.0002). d) fractional change in ischemic lesion volume, e) ^1^H signal within region of interest corresponding to ischemic lesion on day 1 superimposed on subsequent days, f) fractional change in signal (Day 21 control to healthy p = 0.0300) and g) ^1^H signal in contralateral (non-ischemic) hemisphere between groups (* Day 7: p = 0.0094) and (ǂ) naïve animals. Graphs are depicted as mean with standard deviation error bars. Significance was determined by mixed-model with Student’s T test with p < 0.05.

Although providing insight into ischemic lesion volume, traditional ^1^H MRI is mostly based on water content and does not provide direct information on cell membrane integrity or metabolic response. A more sensitive metric, such as ^23^Na CSI, is needed to discriminate therapeutic potential between donors.

### Neuronal and energetic markers differentiate treatment response in stroke

To investigate tissue recovery *in vivo* on a metabolic level, ultra-high field RE-MRS was used to assess lactate (Lac), creatine (Cre) and N-acetyl aspartate (NAA) compared to choline (Cho) levels. Metabolite levels were evaluated in both ischemic and contralateral hemispheres, and were compared to naïve animals (**Fig 4**).

**Fig. 4.**
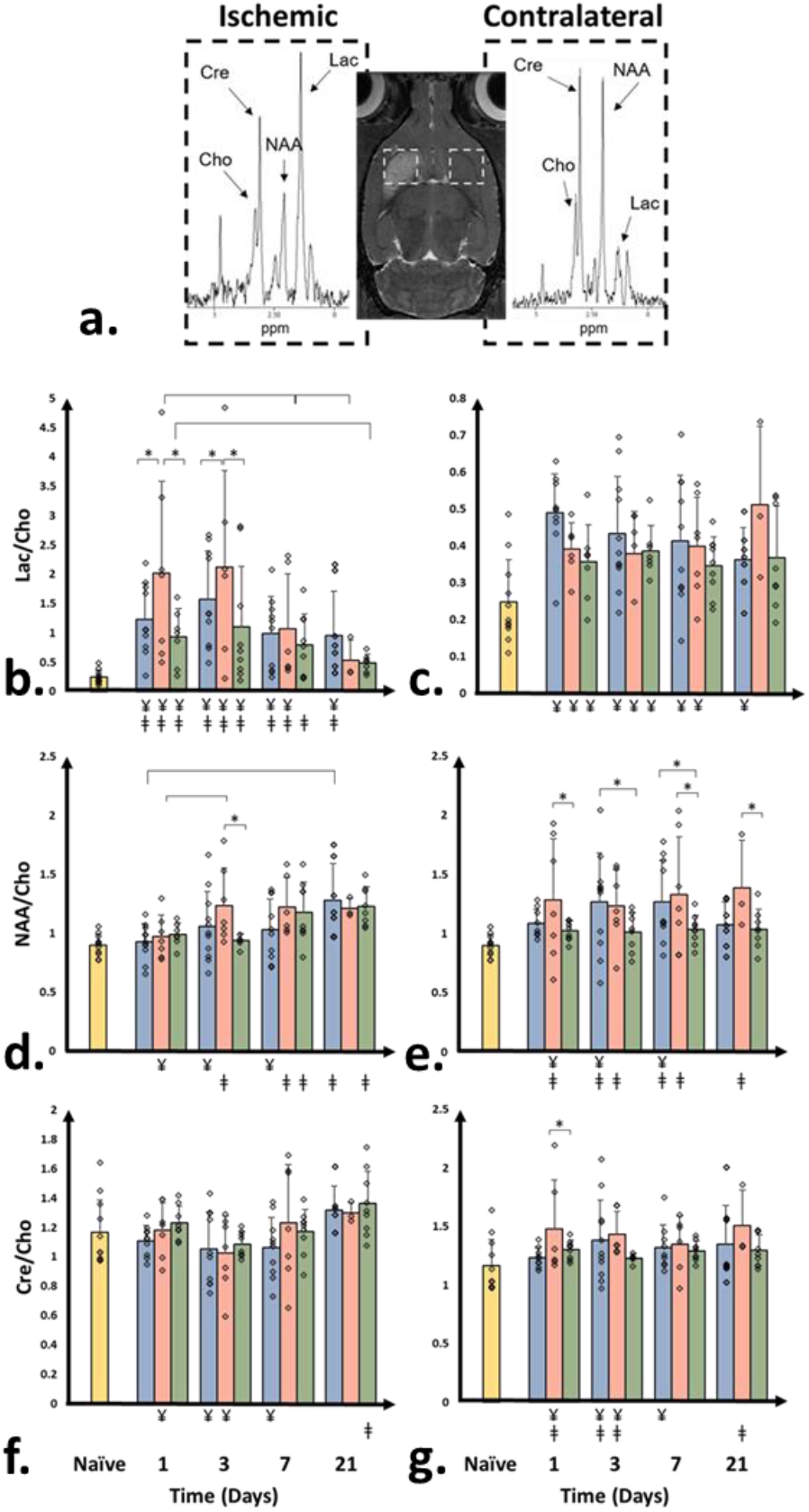
Metabolite levels indicate immediate energetic recovery in the group administered healthy hMSC. a) Representative spectra in ischemic and contralateral hemispheres 1 d after MCAO with localization referenced to anatomical ^1^H T_2_W image. All data sets compared longitudinally, between (*) groups, to (ǂ) naïve levels and between (¥) hemispheres. Lac/Cho levels in b) ischemic (* Day 1: control to compromised p = 0.0023; compromised to healthy p = 0.0001. Day 3: control to compromised p = 0.0425; compromised to healthy p = 0.0003) and c) contralateral hemisphere. NAA/Cho levels in d) ischemic (* Day 3: compromised to healthy p = 0.0311) and e) contralateral hemisphere (* Day 1: compromised to healthy p = 0.038. Day 3: control to healthy p = 0.0258. Day 7: control to healthy p = 0.0365; compromised to healthy p = 0.0177. Day 21: compromised to healthy p = 0.0361). Cre/Cho levels in f) ischemic and g) contralateral hemisphere (* Day 1: compromised to healthy p = 0.0329). Graphs are depicted as mean with standard deviation error bars. Significance was determined by mixed-model with Student’s T test with p < 0.05.

An initial time point, as early as 24 h, was established for distinguishing an effective therapy via Lac/Cho levels. Although Lac/Cho levels were elevated in the ischemic region of all animals at 24 h (p < 0.005), the healthy group demonstrated reduced Lac/Cho at this time compared to the compromised group and maintained this significance even at 72 h for both groups, following a similar trend seen with sodium levels (**Fig. 4b**). Lactate levels returned to naïve baseline levels for the healthy group by day 7 while the compromised group only reached baseline on day 21, and control animals maintained elevated levels throughout the time course. In addition, healthy cells exhibited comparable levels across hemispheres at later time points contrary to the control and compromised groups. Elevated contralateral hemisphere Lac/Cho levels were compared to naïve animals but did not exhibit significance (**Fig. 4c**). Absolute quantification of metabolites normalized to naïve counterparts as a ratio of ischemic region to contralateral hemisphere also were evaluated [**Fig. S1**]. Lactate levels demonstrated significantly increased levels for compromised hMSC compared to control and healthy on days 1 and 3.

Lac/Cho ratios may improve prediction of stroke outcome in clinical settings when combined with other MR metrics [38]. Increased lactate levels occurs as a result of anaerobic glycolysis, the predominant metabolism pathway following an ischemic event [16]. The resulting acidosis, with pH decreasing to 6.4-6.0, impairs mitochondrial respiration, and importantly, neuronal acid-sensing ion channels become activated [39]. This non-selective cation channel is important for mediating the Na^+^ influx, which has been shown here and previously to be a sensitive metric for initial ischemic insult and tissue recovery. This phenomenon also has been studied in cardiac ischemia events [40]. Inflammatory cells, particularly macrophages, have been associated with anaerobic glycolysis, and typically appear three days after initial cerebral ischemic event [41]. The recruitment of macrophages could support the continuous rise in Lac/Cho levels for the control group and maintenance of elevated levels for compromised group, particularly if the elevated Lac/Cho ratio on day 1 is reflective of early inflammatory cell infiltration. No elevation in Lac/Cho levels was evident with a successful therapeutic hMSC therapy onboard, indicating the initial restoration of metabolic process concurrent with reduced inflammatory infiltration.

Typically, NAA initially decreases in ischemic tissue compared to the contralateral hemisphere [42], as it does in this study [**Fig. S1**]. The healthy group maintained baseline levels of NAA/Cho in both the ischemic (**Fig. 4d**) and contralateral (**Fig. 4e**) hemispheres, particularly at early time points. Interestingly, by day 3, NAA/Cho increased for the compromised group compared to healthy. Previous histological studies have demonstrated that, immediately following ischemic damage, NAA may remain trapped in nonviable neurons, which develop into neuronal debris and release NAA into extracellular spaces [43]. Dialysate NAA concentrations, measured via microdialysis and indicative of interstitial NAA, increase following cerebral ischemia [43,44]. Although this increase represents only 2-6% of total cerebral NAA [45], RE-MRS is weighted towards measurement of metabolites with long T_2_ and could correspond to a more mobile NAA population in the interstitial space rather than strictly intracellular NAA. These dynamics could explain the early higher NAA/Cho levels evident for the compromised group on day 3. This loss of neuronal density parallels the concurrent increase in lactate seen in the ischemic hemisphere.

Although a marker for neuronal density, NAA also is involved in the metabolic activity of neurons. Contralateral hemisphere evaluation manifested in robust differences between groups, possibly reflecting compensatory metabolic activity. Most notably, increases in NAA/Cho reflect enhanced mitochondrial activity within neurons for the compromised group over the entire course and the control group on days 3 and 7, while the healthy hMSC group maintained naïve baseline levels throughout (**Fig. 4e**). The contralateral increases in NAA/Cho compared to naïve may represent forced metabolic remodeling in response to systemic toxicity for both the control and compromised groups, in agreement with absence of lactate increases contralaterally.

As an important marker for cell energetics due to its role in generating ATP, creatine also acts as a neuroprotector during oxidative stress [46,47]. Although Cre/Cho levels in the ischemic region (**Fig. 4f**) exhibited comparable levels to naïve baseline for all groups, hemispherical differences were evident early for the compromised group (day 1 and 3) and control animals (day 3 and 7). Within the contralateral hemisphere (**Fig. 4g**), Cre/Cho levels were elevated for the compromised group compared to healthy on day 1 and compared to naïve out to day 21. Typically, creatine depletion is an indicator of cell loss with regard to neural density without specificity to neurons or glia [48]. All groups show a slight decrease in creatine levels in the ischemic hemisphere compared to contralateral, which recovers by day 7 [**Suppl. Fig. 1**]. Cre/Cho ratios may not be as sensitive a metabolic metric as Lac/Cho in assessing longitudinal impacts of cell therapies.

### Behavioral assessment supports hMSC efficacy

To confirm the novel MR approach for assessing hMSC variability, animal weights were monitored and behavior metrics focused on lateral locomotion, anxiety and asymmetry were performed. Animals receiving compromised hMSC demonstrated severe weight loss with delayed recovery, compared to control and healthy hMSC groups (**Fig. 5a**). All behavioral calculations were normalized to individual baseline. The open field test measured lateral locomotion comparing total time spent in the center of the apparatus (**Fig. 5b-d**) as well as the total distance traveled [**Suppl. Fig. 2**] as a test for anxiety. Although only the control group exhibited recovery in lateral locomotion towards baseline (score = 1) from day 3 to days 7 and 14, those receiving healthy hMSC immediately improved in time spent in the center indicating return to normal anxiety levels (score = 1). Asymmetry scores also were evaluated using two methods, an elevated body swing test (**Fig. 5e-g**) and cylinder test [**Suppl. Fig. 2**] with symmetrical score equal to zero. The compromised group maintained an elevated level of asymmetry compared to control on day 21 according to the elevated body swing test. The cylinder test, in which rats voluntarily rear in a clear cylinder apparatus, exhibited reduced symmetry in all groups following ischemia as expected. The compromised hMSC group exhibited increased asymmetry on day 3 compared to control. Increased asymmetry levels have been correlated to decreased recovery following MCAO [49].

**Fig. 5.**
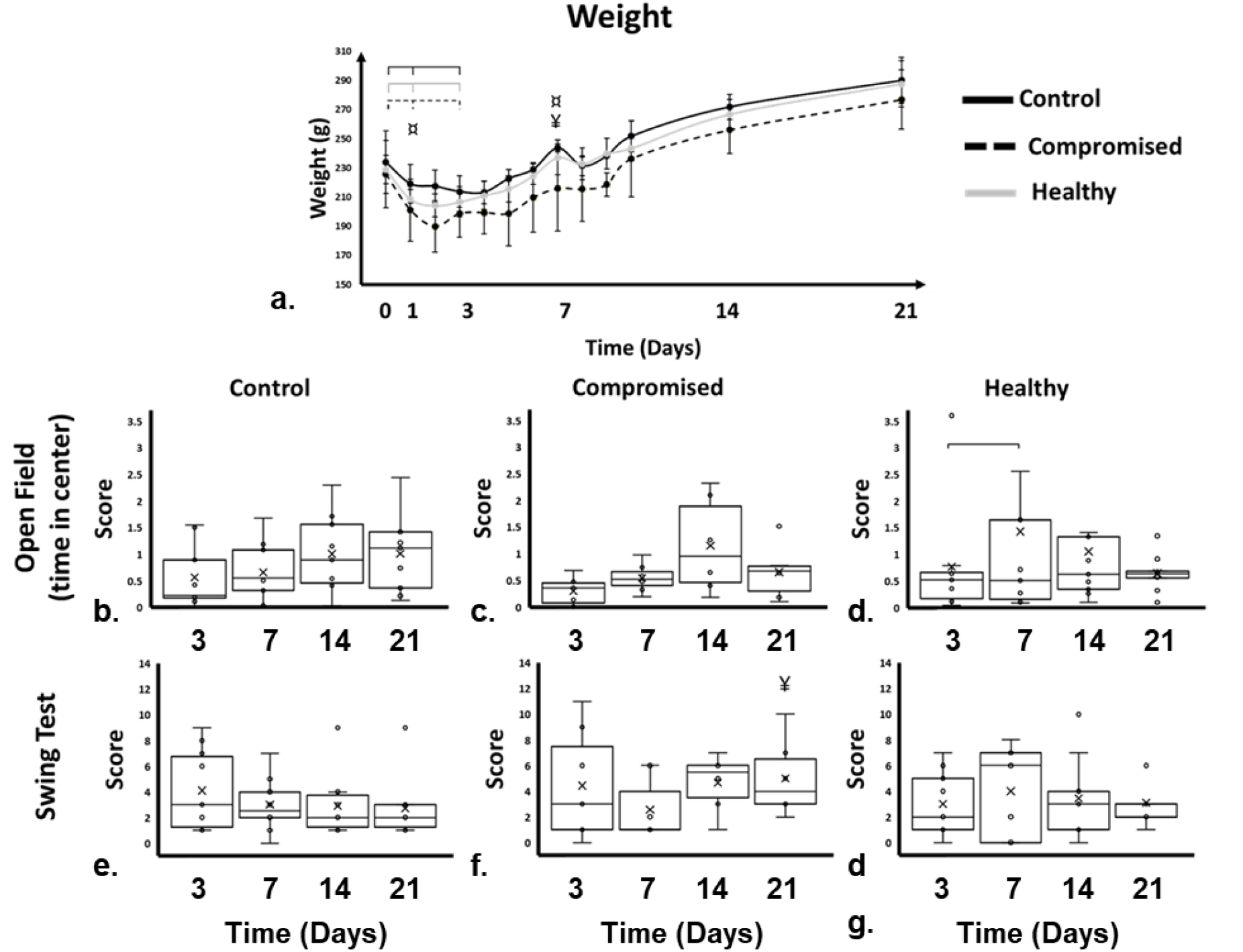
Behavior assessment a) weight in grams of all animals over 21 d time-course. Day 0 to day 1 (control p = 0.0487, compromised p = 0.0051, healthy p = 0.0204) and day 3 (control p = 0.0076, compromised p = 0.003, healthy p = 0.0115). Group comparisons between compromised hMSC and (¤) control (day 1 p = 0.269; day 7 p = 0.0025) and to (¥) healthy hMSC (day 7 p = 0.0234). Time spent in center of open field for b) control, c) compromised hMSC group and d) healthy hMSC group (bar, p = 0.0273). Normalized asymmetry as measured via the swing test for e) control, f) compromised hMSC group (¥, p = 0.0274), and g) healthy hMSC group. Graphs are depicted as mean with standard deviation error bars. Significance was determined by mixed-model with Student’s T test with p < 0.05.

### *In vitro* evaluation corresponding to the time of transplantation cannot determine hMSC deficiency

Hallmarks of cellular senescence or aging are characterized generally by cell proliferation, metabolic activity, SA-β-Gal expression and autophagic potentials [50,51]. The experimental hMSC (both compromised and healthy) were evaluated by these senescent hallmarks and others utilizing various *in vitro* assays. hMSC were tested at passage 5 (P5), corresponding to the point of implantation, as well as following extended culture at P8. At P5, cell morphology (**Fig. 6a**) were similar for both sets of hMSC depicting small, spindle-like structures expected at early passage [52]. In addition, cell proliferation rate characterized by population doubling time (**Fig. 6b**) and colony forming units (**Fig. 6c**) were in line with expected outcomes for MSC at early passage [52]. At early passage, healthy hMCS indicate slightly higher metabolic activity, shifting more strongly towards the oxidative pathway at late passage not reflected with the compromised hMSC [53] [**Suppl. Fig. 3**]. Interestingly, SA-β-Gal activity was comparable at P5 (**Fig. 6d**) indicating the level of senescence in both hMSC groups were similar and therefore qualified for *in vivo* study. Only at extended culture did cell proliferation rate, metabolic activity and SA-β-Gal significantly diverge.

**Fig. 6.**
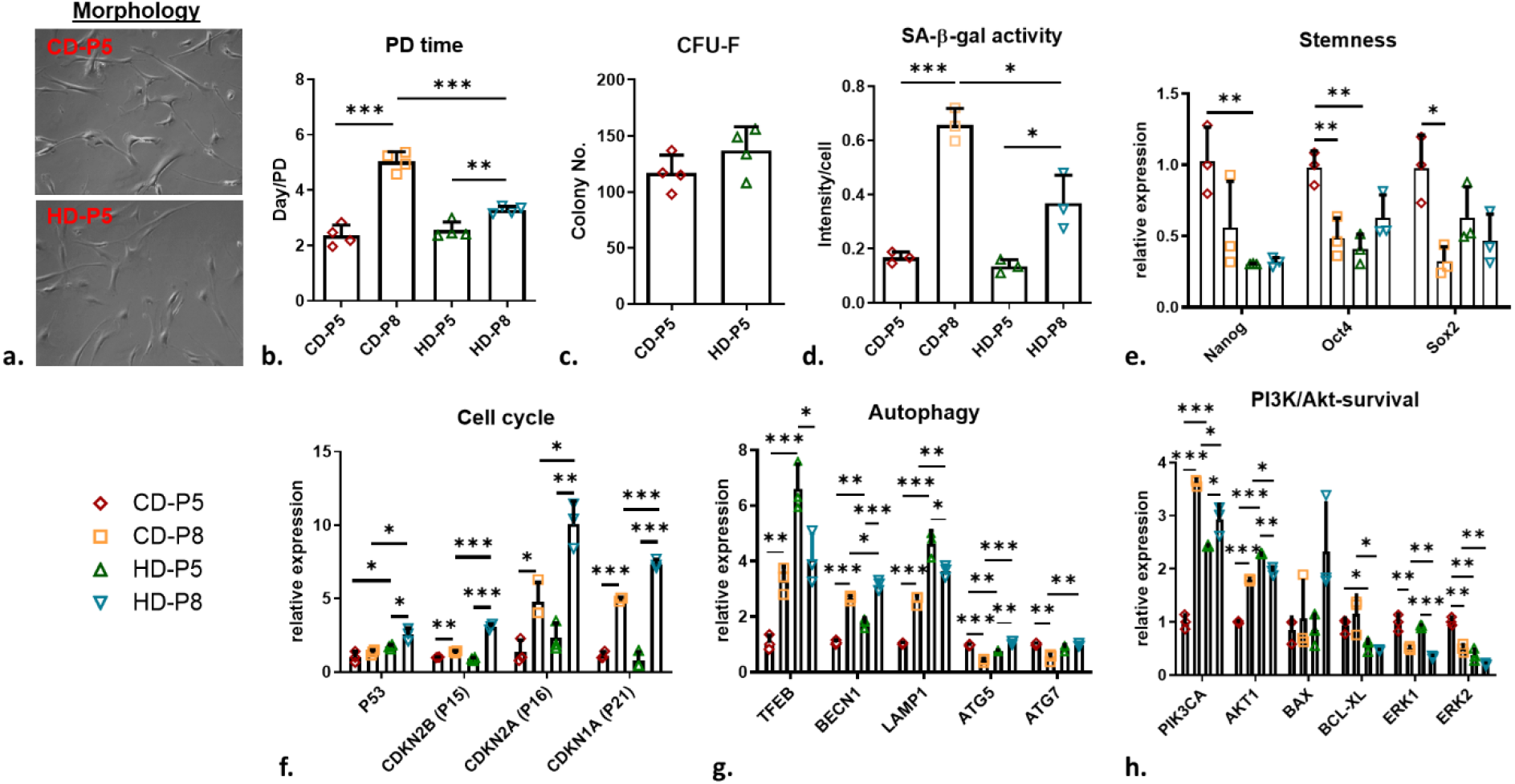
Comparable functionality of hMSC at early passage, corresponding to point of administration is supported by i*n vitro* cell assays. a) Morphology of hMSC at P5 under confocal microscopy. Cell assays for b) population doubling time, c) colony forming, and d) early senescence. RT-PCR for e) stemness, f) cell cycle, g) autophagy, and h) survival at early and late passage. Graphs are depicted as mean with standard deviation error bars and significance was determined by unpaired t test. Significance is marked with * p < 0.05, ** p < 0.01 and *** p < 0.001.

Similarly, differences in stemness or pluripotency gene expressions between healthy and compromised hMSC were most evident in comparison between P5 and P8. Interestingly, Nanog and Oct4 actually displayed significantly higher expression in the compromised hMSC compared to healthy at P5, with Sox2 also elevated (**Fig. 6e**). However, after *in vitro* passage, compromised hMSC displayed decreases in all stem cell genes, significantly so for Oct4 and Sox2, while healthy hMSC expression remained constant between P5 and P8 (**Fig. 6e**). This finding indicates that compromised hMSC were unable to maintain stem cell integrity under culture expansion. Cell cycle genes also supported comparable cell growth at P5 (genes P15, P16 and P21), which became significantly different at P8 (genes P53, P15, P16 and P21) (**Fig. 6f**). Importantly, several genes involved in autophagy displayed significant differences between groups at both P5 and P8 (**Fig. 6g**), which could be a potential checkpoint for evaluation of cell quality [52,54]. In addition, the PI3K/Akt survival pathway, which has key roles for ischemic resistance [55], indicated decreased expression for PIK3CA and AKT1 at early passage (**Fig. 6h**). In contrast to healthy hMSC, severe replicative senescence and reduced stem cell function became evident for compromised hMSC upon *in vitro* culture expansion. The underlying mechanisms that manifest these differential effects with expansion must be present at the low passage used for hMSC transplantation, and would appear to compromise therapeutic potential or even worsen outcomes for disease recovery.

### Additional *in vitro* stress tests confirm compromised efficacy

To evaluate the mechanism potentially at play upon transplantation in compromised hMSC that display replicative senescence, the immunomodulatory potentials, as well as the neuroprotective effects under oxygen-glucose deprivation (OGD), were investigated *in vitro*. Although not significantly different at P5, IDO gene expression was significantly upregulated for both healthy and compromised hMSC by P8 (**Fig. 7a**). However, healthy hMSC IDO gene expression was a factor of 2.2 greater than that seen with compromised hMSC, indicating a higher potential for a self-activated immunosuppressive phenotype after culture expansion. The immunomodulatory potential also was evaluated by IDO activity in each group under IFN-γ priming. Healthy hMSC exhibited higher fold potentials at both P5 and P8 (**Fig. 7b**) with increased IDO activity established when exposed to IFN-γ. The enzymatic activity of IDO is not only involved directly in the inhibition of T cell proliferation but also leads to monocyte differentiation to the more immunosuppressive M2 macrophage phenotype [56].

**Fig. 7.**
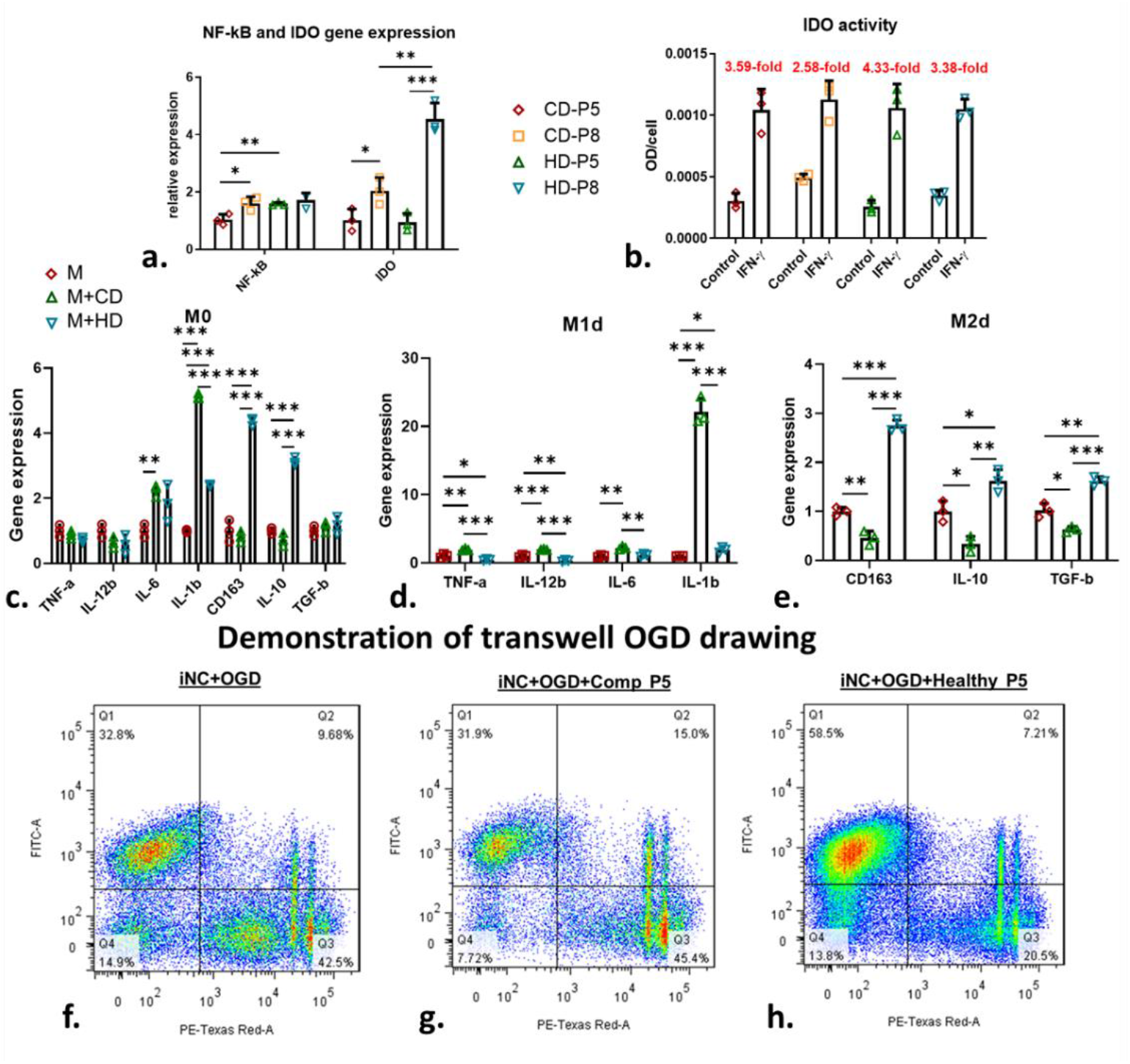
Stress tests differentiate hMSC function further supporting *in vivo* results. a) Gene expression of NF-kB and IDO in compromised and healthy hMSC at P5 and P8. b) IDO activity of compromised and healthy hMSC at P5 and P8 under IFN-y priming. c-e) Macrophage polarization with compromised and healthy hMSC at P5. Genes involved in M1 and M2 phenotype were determined by RT-PCR. f-h) Live/Dead assay of iNC treated with OGD, under co-culture with compromised and healthy hMSC at P5, analyzed via flow cytometry. Graphs are depicted as mean with standard deviation error bars and significance was determine by unpaired T test. Significance is marked with * p < 0.05, ** p < 0.01 and *** p < 0.001

In order to evaluate the role of macrophage involvement and influence from healthy or compromised hMSC further, monocytes differentiated to M0, M1 and M2 macrophages were co-cultured with healthy or compromised hMSC. Compromised hMSC at P5 facilitated the macrophage polarization towards M1 phenotype (**Fig. 7d**) while healthy hMSC improved M2 polarization of macrophages (**Fig. 7e**). Macrophage involvement and recruitment is a leading defense mechanism in brain injuries with the route of activation playing an important role in the innate immune response [57,58]. Classically activated M1 macrophages are associated with the release of pro-inflammatory factors in contrast to the neuroprotective, alternatively activated, M2 macrophages [56,57]. Thus, compromised hMSC may induce the stimulation of pro-inflammatory macrophages in stroke lesion rather than suppress the inflammatory response. This inflammatory response is further supported by the increase in ischemic lesion volume as well as the substantial increase in lactate levels over the first three days following the induction of cerebral ischemia that is seen only in those animals administered compromised hMSC. Alternatively, healthy hMSC induced M2 macrophage polarization may be better suited to influence positively the innate immune response to ischemia in the animal model, which is further supported by the immediate reduction of lesion size and lactate levels.

To test the neuroprotective effect of hMSC, iPSC-derived neuronal cells (iNC) were cultured under OGD condition to mimic the ischemic lesion environment (**Fig. 7f**), and then co-cultured with hMSC via a transwell system. At P5, healthy hMSC improved the neuron survival after OGD (**Fig. 7g**) while the compromised hMSC could not rescue damaged iNC (**Fig. 7h**). Subsequent expansion to P12 further underscored the negative impacts of the compromised hMSC on the iNC culture preparation under OGD [**Suppl. Fig. 4**]. Although it is understood that local response in cerebral ischemia initially favors M2 macrophage phenotype before transitioning to M1, other *in vitro* studies have demonstrated increased oxygen glucose deprivation induced neuronal loss under M1 [57]. As is the case here, compromised hMSC favor M1 macrophage polarization, and thus, the response to OGD is understandably inducing iNC loss. The induction of the M2 alternate by the healthy hMSC protects neurons *in vitro* against OGD, with strong evidence of similar neuroprotection imparted *in vivo* [57]. Ultimately, the compromised hMSC failed to preserve therapeutic potency via *in vitro* tests, which supports pro-inflammatory and low neuroprotective effects that contributed to the decline of therapeutic outcomes in the *in vivo* animal model.

## Discussion

In an ischemic stroke model, 3D ^23^Na MRI and ^1^H MRS are able to differentiate between successful therapeutic agents that otherwise were undiscernible with general *in vitro* testing up to the passage of administration. Only after extended passages and additional stress tests were the hMSC discernable as compromised and healthy, highlighting the need for full characterization of the cells beyond the point of passage at which the cells are injected. These results also support more extensive characterization, including *in vivo* monitoring, to determine whether specific hMSC line or sources are therapeutically beneficial. Thus, a multinuclear MRI/S toolbox for monitoring therapeutic agents *in vivo* offers important and quantifiable metrics to facilitate translational applications. High-resolution 3D ^23^Na MRI quantified restoration of sodium homeostasis, revealing an immediate decrease in lesion volume and signal, 1 to 3 d post-MCAO. This sodium-based assessment was easily distinguishable between treatment groups, providing a previously unseen sensitivity. Importantly, this sensitivity distinguished stem cell treatment efficacy, revealing compromised and healthy cells—a needed step towards useable translational therapies. Energetic recovery of the ischemic region also was demonstrated using ^1^H MRS. Significantly reduced lactate levels immediately following MCAO were observed with accompanying recovery over the time course.

MR findings suggest neuroprotective effects from an optimal therapeutic agent is not limited to the ischemic lesion but extends into the contralateral hemisphere. The contralateral hemisphere was affected both in terms of initial ischemic damage but also with an impactful therapy onboard, as shown by the ability to maintain homeostasis. A level of contralateral metabolic remodeling is suggested primarily in untreated or compromised groups. MRI/S assessment of treatment efficacy mirrors cell assay results that only reveal the difference between donors after extended culture passages or more extensive *in vitro* analysis.

Clinical use of ^23^Na MRI and ^1^H MRS has become a viable option to investigating neurological pathologies with the increased availability of 3-T scanners. ^1^H MRS is particularly advantageous as new hardware is not required, and MRS is already implemented in clinical evaluation of several pathologies, most notably in prostate cancer [59]. Here, we provide evidence that ^23^Na MRI and ^1^H MRS are sufficient at distinguishing effective therapeutics and argue for their implementation in more clinical settings as a tool to differentiate successful therapy in ischemic stroke.

## Acknowledgments

This study was funded by National Institutes of Health and the National Institute for Neurological Disorders and Stroke (RO1-NS102395 and F31-NS115409). The content is solely the responsibility of the authors and does not necessarily represent the official views of the National Institutes of Health. A portion of this work was performed at the National High Magnetic Field Laboratory, which is supported by National Science Foundation Cooperative Agreement No. DMR-1644779 and the State of Florida.

## Author Contributions

S.H. and S.C.G. conceived, designed and directed the project. S.H. drafted the manuscript, performed all surgical procedures, and acquired and interpreted all MR data. X.Y. designed and performed all cell culture and *in vitro* analysis. J.A. performed and compiled blind animal behavioral assessments and S.H. interpreted the results. F.A.B. assisted in animal handling and MR acquisition as needed. C.V.B assisted with interpretation. S.C.G. supervised the project. All authors contributed to and approved the manuscript.

## Ethic Statement

All applicable international, national and/or institutional guidelines for the care and use of animals were followed. All animal procedures were completed in accordance with the Animal Care and Use Committee at the Florida State University. All studies were conducted in accordance with the United States Public Health Service’s Policy on Humane Care and Use of Laboratory Animals as well as the US NIH Guide for the Care and Use of Laboratory Animals (NIH Publications, No. 8023, 1978 revision).

## Data Statement

All data used in this study were newly acquired. Relevant data used in preparation of this manuscript are available upon request to either the principal or corresponding author via email. In accordance with the National High Magnetic Field Laboratory FAIR Data Management Plan (www.nationalmaglab.org/about/fair-data), data can be made available for access through the NHMFL publication database.

## Additional Information

Supplementary Information is available for this paper. Correspondence and request for materials should be addressed to Samuel C. Grant.

