## Supplemental Information for "Multinuclear MRI reveals early efficacy of stem cell therapy in stroke"

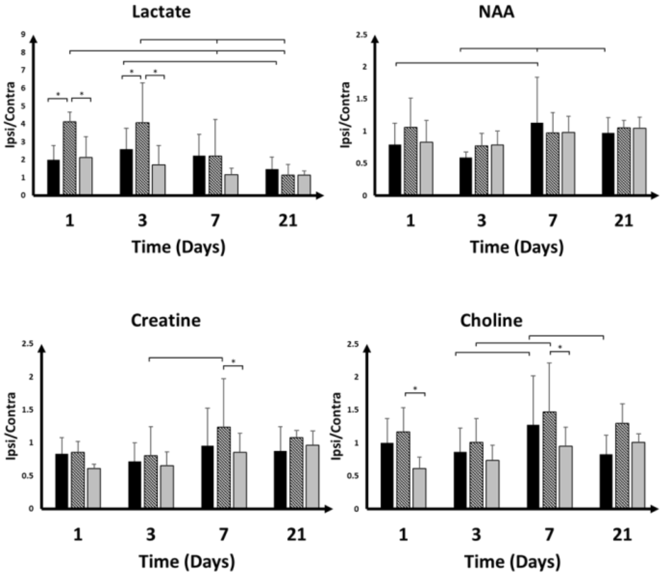


**Figure S1. Absolute quantification of metabolites normalized to naïve counterparts.**

$^{1}$H MRS demonstrates metabolites as a ratio of ischemic to contralateral hemisphere normalized to naïve rats. Lactate levels indicate significantly elevated levels in rats having been administered compromised hMSC compared to control and healthy hMSC. An immediate reduction of lactate from day 1 to 3 is evident for the healthy hMSC group. Graphs are depicted as mean with standard deviation error bars. Significance was determined by mixed-model with Student's T test with $p < 0.05$.

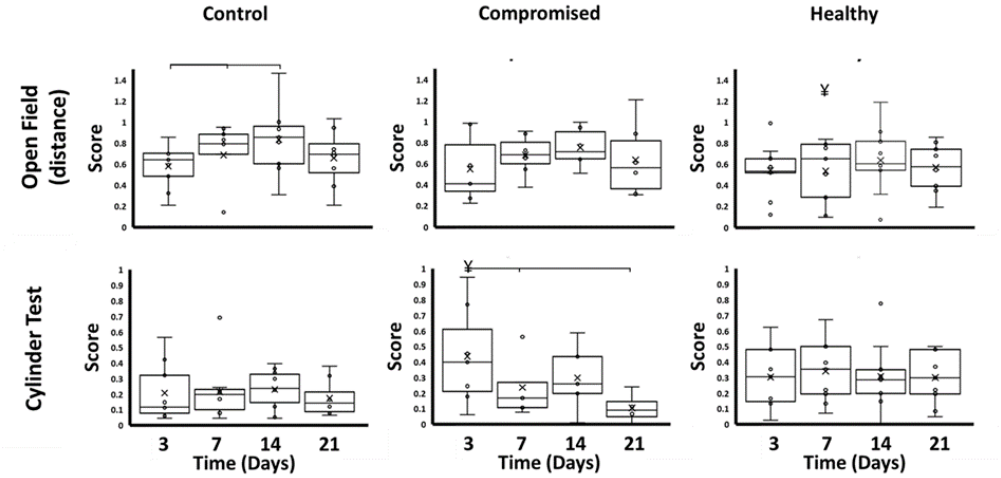


**Figure S2. Behavioral characteristics**

Distance travelled in open field test is demonstrated for all three groups over 21 d. Cylinder test represents increased asymmetry on day 3 for compromised group only. Graphs are depicted as mean with standard deviation error bars. Significance was determined by mixed-model with Student's T test with $p < 0.05$.

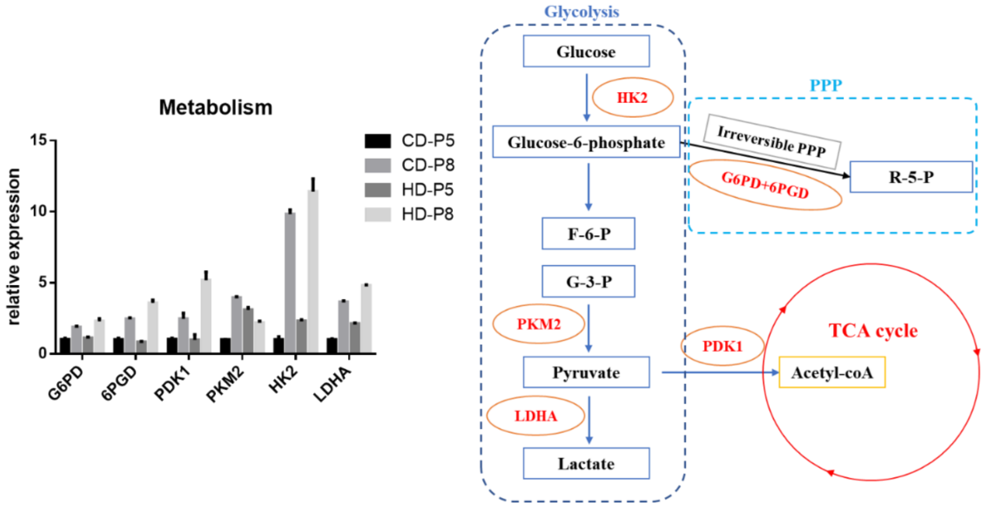


**Figure S3. Metabolism**

Relative gene expression indicative of metabolic pathways for compromised and healthy hMSC with visual schematic.

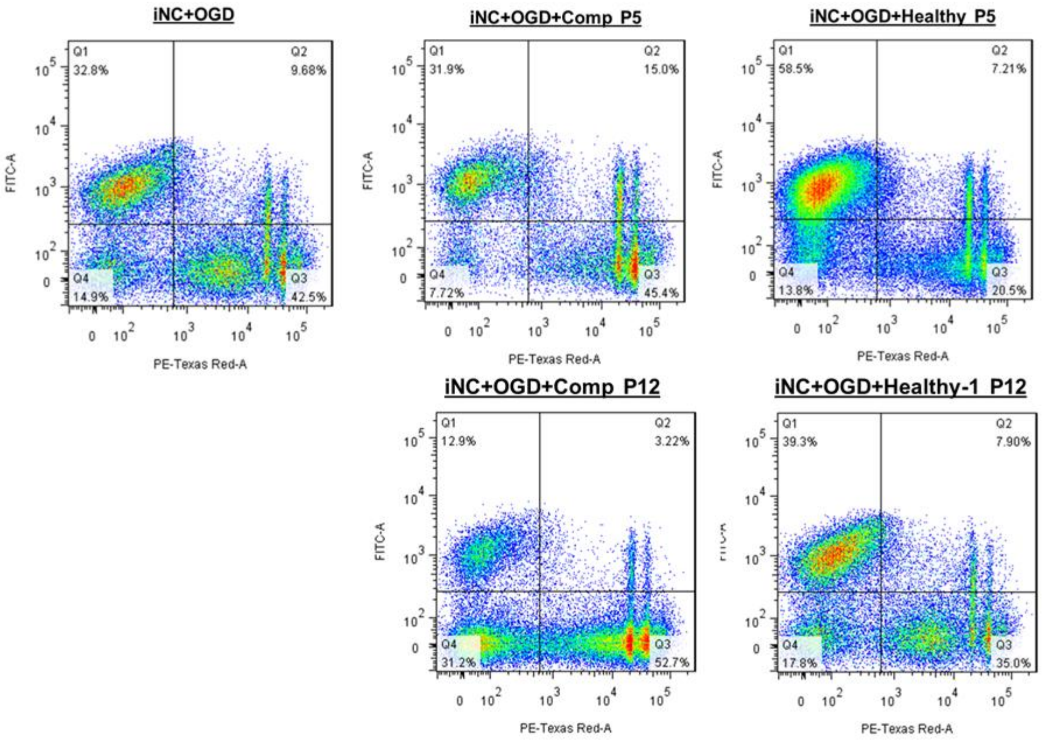


**Figure S4. Oxygen-Glucose Deprivation test**

To test the neuroprotective effect of hMSC, iPSC-derived neuronal cells (iNC) were cultured under OGD condition to mimic ischemic lesion environment, and then co-cultured with hMSC via a transwell system. At P5, healthy hMSC improved the neuron survival after OGD while the compromised hMSC could not rescue damaged iNC. Subsequent expansion to P12 further underscored the negative impacts of the compromised hMSC on the iNC culture preparation under OGD.
